# *Fusarium verticillioides* SNARE protein FvSyn1 harbors two key functional motifs that play selective roles in fungal development and virulence

**DOI:** 10.1101/615211

**Authors:** Huan Zhang, Huijuan Yan, Won Bo Shim

**Author notes:** **For Correspondence:** Huan Zhang.

## Abstract

*Fusarium verticillioides* is one of the key fungal pathogens responsible for maize stalk rots. While stalk rot pathogens are prevalent worldwide, our understanding of stalk rot virulence mechanism in pathogenic fungi is still very limited. We previously identified *F. verticillioides FvSYN1* gene, which was demonstrated to plays an important role in maize stalk rot virulence. FvSyn1 belongs to a family of soluble *N*-ethylmaleimide-sensitive factor attachment protein receptor (SNARE) proteins that play critical roles in a variety of developmental processes. In this study, we further characterized the cellular features of FvSyn1 protein motifs in *F. verticillioides* development and virulence. We generated FvSyn1 motif-specific deletion mutants to further investigate how different motifs contribute to development and virulence. Microscopic observation showed that Δfvsyn1 mutant exhibits rough and hyper-branched hyphae when compared to the wild type progenitor. Moreover, Δfvsyn1 mutant was sensitive to cell wall stress agents resulting in vegetative growth reduction. We showed that FvSyn1::GFP protein is associated with endomembrane but this outcome did not clarify why the deletion of this protein led to stress sensitivity and aberrant hyphal development. Characterization of FvSyn1 domains indicated that both Syntaxin N-terminus (SynN) domain and SNARE C-terminus domain play distinct roles in fungal development but collectively function in virulence. We also determined that two domains in FvSyn1 is not required for fumonisin production. Interestingly, these two domains were involved in carbon nutrient utilization including pectin, starch and sorbitol. This study further characterized the role of FvSyn1 in hyphal growth, localization, cell wall stress response and virulence in *F. verticillioides*.

**Highlights:** ► *F. verticillioides* SNARE protein *FvSYN1* is important for vegetative growth and virulence.

► *FvSYN1* deletion mutant performed better under cell wall stressors.

► Dissecting of two domains and investigate the roles.

## 1. Introduction

Stalk rot is one of the most persistent diseases of maize, which is primarily caused by a group of fungal pathogens. *Fusarium verticillioides, F. graminearum*, *Colletotrichum graminicola* and *Macrophomina phaseolina* are the most common stalk rot pathogens that are found in maize-growing regions in the US (Koehler, 1960; Kommendahal & Windels, 1981; White, 1999). Stalk rot pathogens typically overwinters in crop residue by producing thickened hyphae and spores (Leslie *et al*., 1990). Stalk rot pathogens are commonly known to penetrate maize through natural entry points or wounds created by insects or mechanical damages at the base of leaf sheaths and progress to lower internode (Foley, 1962; Kingsland & Wernham, 1962; Koehler, 1960; Sobek and Munkvold, 1999). It is also suggested that the disease is seed-borne or seed-transmitted (McGe, 1988). Pathogen invasion and colonization prior to physical maturity of crops usually result in yield reduction. Abiotic stress factors, particularly near the end of growing season when the developing ear competes with the stalk for nutrients, are also known to have a major impact on stalk rot outbreaks (Dodd, 1980; Smith &White, 1989).

We currently have very limited understanding of molecular genetic mechanism associated with fungal virulence in this complex disease. One of the few stalk rot virulence genes characterized is *F. verticillioides FSR1*, which encodes a protein similar to mammalian striatin (Shim *et al*., 2006; Yamaura & Shim, 2008). Striatin is known to form a large multi-protein assembly referred to as the striatin-interacting phosphatase and kinase (STRIPAK) complex, and this complex has also been reported in filamentous fungi (Beier *et al*., 2016; Dettmann *et al*., 2013). Our recent study further determined that Fsr1 directly interacts with three newly identified proteins not associated with STRIPAK to regulate stalk rot virulence (Zhang *et al*., 2018). Subsequently, we aimed to further characterize transcriptional networks downstream of Fsr1 by performing next-generation sequencing (NGS) with maize B73 stalks inoculated with *F. verticillioides* wild type and *fsr1* mutant. We established a computational analysis pipeline, including probabilistic pathway activity inference method, to identify functional subnetwork modules likely involved in *F. verticillioides* virulence (Kim *et al*., 2015; Kim *et al*., 2018a). Through our analyses, we identified a putative virulence gene *FvSYN1* from virulence-associated subnetwork modules (Kim *et al*., 2018b). FvSyn1 is predicted to encode a soluble *N*-ethylmaleimide-sensitive factor attachment protein receptor (SNARE) with two well-recognized functional domains including a Syntaxin N-terminus (SynN) domain and a SNARE C-terminal domain.

SNARE proteins play critical and conserved roles in eukaryotic intracellular membrane fusion during various trafficking processes, notably in the secretory and endocytic pathways (Chen & Scheller, 2001; Hong & Lev, 2014). There are diverse sets of SNARE proteins found in different cellular compartments, with each SNARE protein forming a specific SNARE complex with different membrane trafficking steps (Ryu *et al*., 2016; Scales *et al*., 2000). Despite the diversity, all SNARE proteins contain a conserved SNARE domain that consists of approximately 60-70 amino acids with multiple coiled-coil structures forming heptad repeats in the membrane-proximal regions of the SNARE proteins (Chen & Scheller, 2001; Hong & Lev, 2014). Moreover, many SNAREs contain N-terminal regulatory region (Hong & Lev, 2014). SNAREs were originally classified into v-SNAREs and t-SNAREs according to their vesicle or target membrane localization (Söllner *et al*., 1993). However, many SNAREs were later found both on vesicles and target membranes, and SNAREs were reclassified into Q-SNAREs (glutamine-containing SNAREs) and R-SNAREs (arginine-containing SNAREs) according to the crystal structures of the synaptic SNARE complex (Fasshauer *et al*., 1998). R-SNAREs act as v-SNAREs, and most Q-SNAREs act as t-SNAREs.

SNARE proteins are well characterized in mammals, and the known mammalian SNAREs have been classified into syntaxin, VAMP (also called synaptobrevin), and SNAP25 families. Recent studies also reported that SNARE proteins are important for vesicle fusion and development in yeast and filamentous fungi (Gupta & Heath, 2002). In *Saccharomyces cerevisiae*, SNARE Sso1 and Sso2 exhibited functional redundancy in affecting vesicle fusion and sporulation (Nakanishi *et al*., 2006). In *Neurospora crassa*, *nsyn1* (Sso1 homolog) was shown to be important for hyphal development, conidiation and male fertility while *nsyn2* (Sso2 homolog) was important for hyphal branching and ascospore development (Gupta *et al*., 2003). In wheat scab pathogen *F. graminearum*, Hong *et al*. (2010) reported *GzSYN1* and *GzSYN2* genes, with *GzSYN1* deletion mutant showing reduced hyphal growth rate while *GzSYN2* deletion mutant exhibiting fertility issues. Both genes were shown to be important for virulence in this pathogen (Hong *et al*., 2010). Our previous study showed that *FvSYN1*, a downstream gene of *FSR1*, plays important roles in vegetative growth and maize seedling rot (Kim *et al*., 2018b). The main objectives of this study were to further characterize the roles of FvSyn1 in *F. verticillioides* to gain deeper understanding of stalk rot pathogenesis. We studied vegetative growth and hyphal development, as well as the cellular localization of the protein. We also investigated the functional roles of two well-recognized domains, N-terminal SynN domain and C-terminal SNARE domain, in FvSyn1.

## 2. Material and methods

### 2.1 Fungal strains, culture media and nucleic acid manipulation

*F. verticillioides* wild-type strains (7600 and 7598) and all mutants used in this study were cultured at 25°C on V8 agar (200 ml of V8 juice, 3 g of CaCO_3_ and 20 g of agar powder per liter) for inoculum preparation and routine maintenance. Colony morphology was visually assayed on V8 agar and potato dextrose agar (PDA; Difco). For stress assay, spores of all strains were cultured on PDA amended with different concentrations of NaCl, KCl, sorbitol, calcofluor white (CFW), sodium dodecyl sulfate (SDS) and congo red (CR), and then incubated at 25 ° for 5 days. Standard molecular manipulations were performed as described previously (Sagaram & Shim, 2007). Strains were grown in YEPD liquid medium (3 g yeast extract, 10 g peptone and 20 g dextrose per liter) at 25 °C with agitation (150 rpm) and harvested for genomic DNA extraction using the OminiPrep genomic DNA extraction kit (G Biosciences, Maryland heights, MO, USA). For carbon source utilization assay, spores of all strains were spotted on the center of the Czapek-Dox agar plate (2 g/L NaNO_3_, 0.5 g/L MgSO_4_·7H_2_O, 0.5 g/L KCl, 10 mg/L 14 FeSO_4_·7H_2_O, and 1 g/L K_2_HPO_4_, pH 7) with different carbon sources (sucrose 30 g/L, pectin 10g/L, sorbitol 182.17/L or starch 15 g/L). Vegetative phenotypes and growth rates (colony diameter) were assayed after 8 days of incubation at room temperature.

### 2.2 Phylogenetic analysis

Putative FvSyn1 orthologs were identified by protein-protein BLAST analyses (BLASTp). Sso1 protein structure was obtained from PDB (PDB ID:1FIO) (Munson *et al*., 2005) and modified into alignment by ESPript (Robert and Gouet, 2014). Multiple sequences were aligned with the ClustalX2 program. For phylogenetic tree generation, the sequences aligned by ClustalX2 were subjected to the tree building process with Neighbour-Joining method by Mega5 with 1000 bootstrap (Tamura *et al*., 2011).

### 2.3 GFP constructs generation

For *in vivo* localization, we generated GFP strains by introducing FvSyn1::GFP fusion construct under the control of the RP27 promoter into the corresponding mutant protoplast as described previously (Zhang *et al*. 2018). GFP was amplified from gGFP using primers sGFP/F and5GAsGFP/Rbs with five glycine–alanine repeat (GA-5) sequences attached at the N-terminus as a linker for GFP tagging at the N-terminus. Primers RP27-F and RP27-R were used to amplify RP27 promoter from PET11 plasmid (Zhang *et al*. 2018). Primers Syn1-GFP-F/R were used to generate *FvSYN1* gene fragment. To generate RP27::GFP::FvSyn1, the RP27::GFP fusion construct was generated first, then fused with *FvSYN1* by joint PCR method. All primers used in this study are listed in Table S1. The construct, together with a geneticin-resistance (*GEN*) marker, were introduced into the Δfvsyn1 mutant protoplasts following the standard *F. verticillioides* transformation protocol (Sagaram & Shim, 2007).

### 2.4 Cytological Assay

To assess hyphal growth and morphology, wild-type (WT), Δfvsyn1 mutant and complementation (Fvsyn1C) strains were first inoculated on V8 plates for 7 days. Subsequently, conidia were collected, inoculated into a 96-well plate which contained 200 μl potato dextrose broth (PDB) and further incubated at 25 °C. Conidia germination was observed under Optika XDS-2 microscope at 0 hour, 9 hours and 27 hours post inoculation. For macrospore observation, strains were inoculated on V8 plates for one week and spores were collected and observed under Olympus BX60 microscope.

To visualize subcellular localization of FvSyn1, the spores of GFP-tagged strains were incubated in PDB for 15-20 hours at room temperature. Conidia, germinated conidia, and vegetative hyphae were observed with inverted Nikon Eclipse-Ti microscope with a 100× 1.49 NA TIRF objective, and the images were taken using a Hamamatsu ImagEM ×2 EMCCD camera C9100-23B. To visualize the cell endomembrane, hyphae were treated with 25 μM FM4- 64 solution for 20 min before being observed under the microscope. For endocytosis assay, hyphae were treated with FM 4-64 for 1 min before observed under microscope. All GFP images were prepared for publication using ADOBE PHOTOSHOP CS5.1 (Adobe).

### 2.5 Generation of FvSyn1 motif-deletion mutants

To investigate the functional role of FvSyn1 N-terminus and C-terminus regions, we first generated two complementation constructs. One has a complete deletion of the N-terminus domain (Δ*SynN*) with native promoter and terminator, and the other has a complete deletion of the C-terminus (Δ*SNARE*) with native promoter and terminator. Both were generated via homologous recombination. For Δ*SynN* construct, primers 1 and 2 were used for SynN 5’ flanking region generation, and primers 3 and 4 were used for SynN 3’ flanking region generation. Both 5’ and 3’ flanking region fragments were then fused by single-joint PCR using primers 1 and 4 (Fig. 5A). Using the same strategy, Δ*SNARE* construct was generated. Then we introduced the constructs together into Δfvsyn1 mutant protoplasts along with a geneticin-resistance (*GEN*) marker. PCR was performed to screen the motif deletion mutants, and qPCR was performed to confirm the mutations (Fig. S2). All primers used in this study are listed in Table S1. Transformants were regenerated and selected on regeneration medium containing 100 µg/ml of hygromycin B (Calbiochem, La Jolla, CA, USA) and 150 µg/ml G418 sulfate (Cellgro, Manassas, VA, USA). Respective drug-resistant colonies were screened by PCR. Complementation strain with complete wild-type gene driven by its native promoter was also generated (Kim *et al*., 2018b) as a positive control to study motif-deletion mutants.

### 2.6 Maize infection assays

Maize seedling rot assay was performed on 2-week old Silver Queen (Burpee) maize seedlings as previously described (Christensen *et al*., 2014; Zhang *et al*., 2018) with minor modifications. Briefly, spore suspensions in 0.1% Tween-20 (1×10^8^/ml) were prepared for maize mesocotyl infection. Plant mesocotyls were first slightly wounded by a sterile syringe needle about 3cm above the soil. A 5μl-spore suspension was applied to the wound site. The seedlings were immediately covered with a plastic sandwich bag to create a high moisture environment suitable for infection and colonization. The seedlings were collected and analyzed by Image J software after a 2-week growth period in the dark room. At least three biological and three technical replicates were performed for each fungal strain.

### 2.7 FB_1_ and ergosterol analysis

FB_1_ analysis was performed following the established protocol (Shim & Woloshuk, 1999) with some modifications. Wild-type and mutant conidia (2×10^6^) were inoculated into 2 g cracked corn, where kernels were placed in 20-ml glass vials, rehydrated in 1.5 ml water overnight and autoclaved for use, for 7 days at 25°C. Acetonitrile: water (10 ml, 1:1, v/v) was added to the vials and incubated at room temperature 24 hours without agitation for FB_1_ extraction. Chloroform:methanol (10 ml, 2:1, v/v) was added to the vials for ergosterol extraction. FB_1_ extracts were purified by passing through equilibrated SPE C18 columns (Fisher Scientific), washed with water, followed by 15% acetonitrile and then eluted with 70% acetonitrile. Ergosterol extracts were passed through acrodisc 13 mm Nylon 0.45-μm filters (Pall Life Sciences). FB_1_ and ergosterol HPLC analyses were performed as previously described (Kim *et al*., 2011; Shim & Woloshuk, 1999). Quantifications of FB1 and ergosterol were done by comparing HPLC peak areas to FB1 and ergosterol standards (Sigma). FB_1_ levels were then normalized to ergosterol contents in samples by calculating [FB_1_ ppm / ergosterol ppm] × 100 (Kim *et al*., 2011).

### 2.8 Sexual cross experiment

*F. verticillioides* sexual cross was performed as described by Sagaram and colleagues (2007). All strains were first grown on V8 plates, and then conidia of the wild-type strain 7598 (*MAT1-2* genotype) were collected and spread to carrot agar plates and incubated for 7 days at 25°C. Then conidia of the wild type 7600 (*MAT1-1* genotype) and mutant strains generated in this background were harvested and quantified, same amount of conidia (5×10^6^ spores) were applied to plates covered with strain 7598 strain. Crosses were maintained at 25 °C until perithecia and ascospores were observed and characterized.

## 3. Results

### 3.1 *F. verticilloides* FvSyn1 is a conserved SNARE protein

*FvSYN1* encodes a putative 375-amino-acid protein with an N-terminus SyN domain and a C-terminus SNARE domain (Fig.1A). To identify the orthologs in other fungal species, FvSyn1 amino acid sequence was used for a BLASTp search. Fvsyn1 protein query returned high identity with its SNARE orthologs in *F. graminearum* GzSyn1 (FGSG_00950, 89%identity), *Magnaporthe oryzae* MoSso1 (MGG_04090, 66%identity), *N. crassa* NSyn1 (NCU02460, 60%identity), *Colletotrichum graminicola* CgSso1 (GLRG_05973, 63%identity) and *Saccharomyces cerevisiae* Sso1 (SCY_5500, 30% identity) (Fig. 1B). Multiple sequence alignment and phylogenetic tree analyses further confirmed that these SNARE proteins are highly conserved in these fungal species (Fig. 1C). We also identified FvSyn1 homologs in other eukaryotes, namely in human fungal pathogens *Cryptococcus neoformans* (CnSyn1, C349_07154, 31% identity) and *Candida albicans* (CaSyn1, MGE_02362, 33% identity), as well as in *Mus musculus* (MmSyn1, mCG_14375, 28% identity), *Oryza sativa* (OsSyn1, SYP121, 23% identity) and Homo sapiens (HsSyn1, STX2, 26% identity). While Fvsyn1 shares relatively lower similarities with SNARE proteins in other eukaryotes, the results strongly suggest that FvSyn1 is a conserved eukaryotic SNARE protein.

**Fig. 1.**
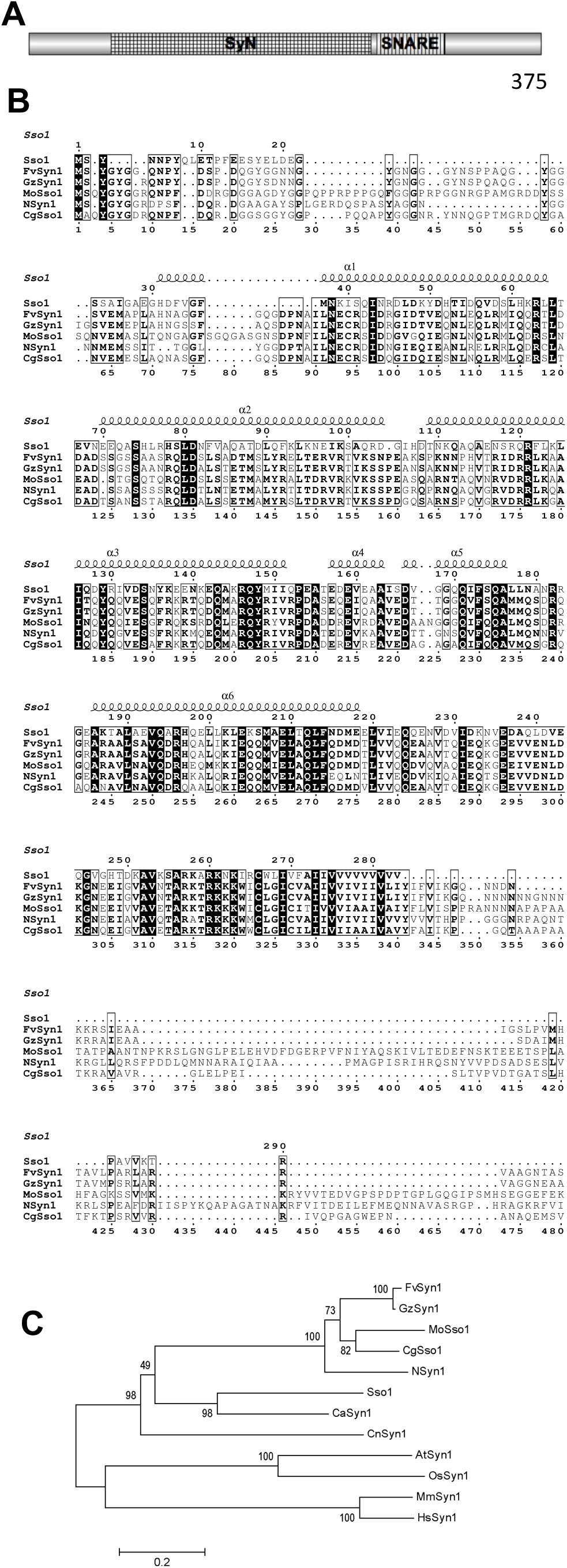
FvSyn1 proteins and its orthologs in other fungal species. (A): Schematic representation of FvSyn1 and its two domains. (**B**): Amino-acid alignment of the amino acids in *F. verticillioides* FvSyn1, *F. gramminum* GzSYN1, *M. oryzae* MoSso1. *C. graminicola* CgSso1, and *S. cerevisiae* Sso1. Conserved amino acids are indicated with a rectangle frame, identical amino acid residues are shaded. (C) Phylogenetic tree for *FvSYN1* gene and its homologs. (D) Maximum likelihood tree for the SNARE gene families. The sequences used in this analysis are *F. verticillioides* FvSyn1 (FvSyn1, FVEG_00594), a *F. gramminum* homolog (FGSG_00950, GzSyn1), a *Magnaporthe oryzae* homolog (MGG_04090, MoSso1), a *Neurospora crassa* homolog (NCU02460, Nsyn1), a *Colletotrichum graminicola* homolog (GLRG_05973, CgSso1), *Saccharomyces cerevisiae* homolog (SCY_5500, Sso1), a *Cryptococcus neoformans* var. neoformans B-3501A homolog (CnSyn1), a *Candida albicans* homolog (CaSyn1), an *Arabidopsis thaliana* homolog (AtSyn1), an *Oryza sativa Japonica* homolog (OsSyn1), a *Mus musculus* homolog (MmSyn1), a *Homo sapiens* homolog (HsSyn1).

### 3.2 FvSYN1 influences hyphal growth and branching

Our previous study (Kim *et al*., 2018b) showed that the Δfvsyn1 mutant exhibited reduced vegetative growth while having more dense and fluffier mycelia on solid agar media when compared with the wild type (WT). But there was no significant difference in fungal mass when we measured the mycelia harvested from YEPD broth. We hypothesized that these outcomes are due to the mutation causing altered hyphal growth and perhaps extensive branching in the Δfvsyn1 strain when cultured under agitation. To test this, we inoculated the spores of WT, Δfvsyn1 and the complementation strain (FvSYN1C) into potato dextrose broth (PDB) and monitored the germination process under a microscope. After incubating for 9 hours, we learned that *FvSYN1* was germinating slower than wild-type and complementation strains (Fig. 2). However, while no significant growth rate difference was observed 27 hours after inoculation, mycelia of Δfvsyn1 exhibited vigorous branching and higher hyphae density when compared to WT and FvSYN1C strains (Fig. 2). These results suggest that *FvSYN1* plays a role in vegetative development and provided explanation for why Δfvsyn1 exhibited slower horizontal vegetative growth on solid media while showing little biomass difference in liquid cultures.

**Fig. 2.**
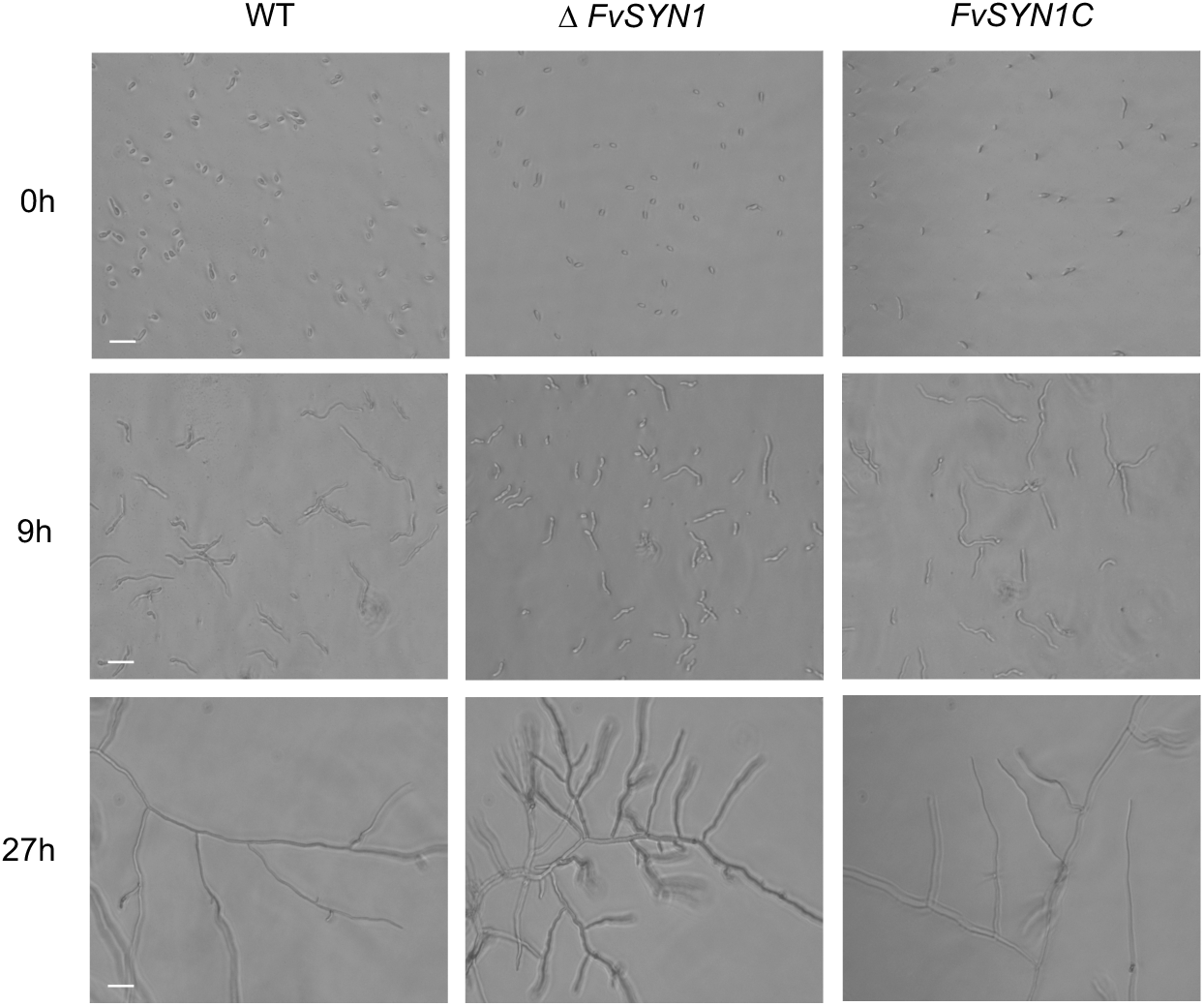
Conidial germination of wild-type (WT), *FvSYN1* gene deletion mutant and complementation strain (*FvSYN1C*) in PDB broth were observed under Optika XDS-2 microscope at different time points. (A-C): Spores of WT, mutant and complementation strains in 96 well plates. (D-F): Spore germination morphology 9 hours after inoculation. (G-I): Hyphae tip area morphology 27 hours after inoculation. Scale bar = 30 μm.

### 3.3 Localization of FvSYN1 in F. verticillioides

*F. verticilloides* FvSyn1 is a conserved SNARE protein and SNARE proteins are known to play roles in various trafficking processes on eukaryotic cellular membranes. We hypothesize that FvSyn1 is likely to have conserved functions in trafficking proteins associated with cellular membranes. We studied the subcellular localization of FvSyn1 in *F. verticillioides* by generating a FvSyn1-GFP fusion construct to transform into Δfvsyn1 mutant. We found that the conidia of FvSyn1-GFP mainly localizes to plasma membrane, the germinating conidia and hyphae were localized to plasma membrane, vacuoles and septa. This was further verified with FM4-64 staining, which preferentially stains endomembrane, which colocalized with FvSyn1-GFP signals (Fig. 3). The result suggested that FvSyn1 is physically associated with endomembrane, but we need to further evaluate whether this SNARE protein is directly involved in protein trafficking.

**Fig. 3.**
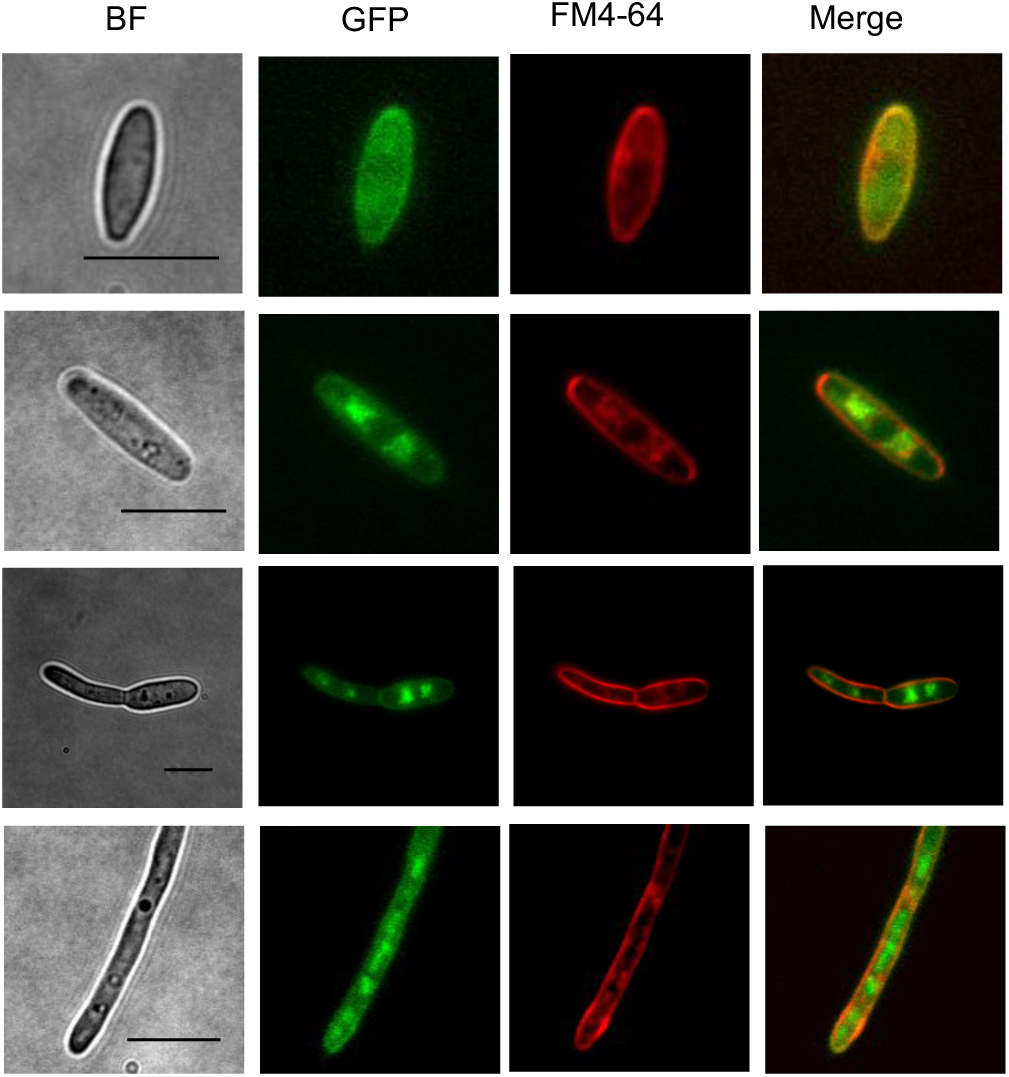
Expression and localization of FvSyn1-GFP in *F. verticillioides*. Spores, germinating spores and vegetative hyphae expressing the FvSyn1-GFP fusion construct were examined. The same field was examined under bright field (BF) and epifluorescence microscopy [green fluorescent protein (GFP) or FM4-64 staining]. Scale bar = 10 μm.

### 3.4 FvSYN1 plays an important role in response to various stressors

To further examine the role of FvSyn1, we tested the growth responses of Δfvsyn1 mutant on solid media containing various cell stressors agents. Strains were inoculated on 0.2×PDA plates amended with KCl (1M), NaCl (1M), sorbitol (1M), sodium dodecylsulfate (SDS) (0.01%), congo red (CR) (100μg/ml) and calcofluor white (CFW) (100μg/ml) (Fig. 4A). As the stressors can inhibit or promote WT growth, we set the growth rate of WT on PDA plates as the standard to normalize other strains and conditions. The mutant growth rates were greater than those of WT and Fvsyn1C when grown on plates with SDS, CR and CFW. However, there were no significant differences when grown on plates with KCl, NaCl and sorbitol (Fig. 4B), although we observed overall growth deficiency on plates amended with KCl and NaCl. Our results suggested that FvSyn1 was involved in controlling cell wall stress response. Similar results in other SNARE proteins such as MoVam*7* in *M. oryzae* and FgVam7 in *F. graminearum* were also reported (Zhang *et al*., 2016).

**Fig. 4.**
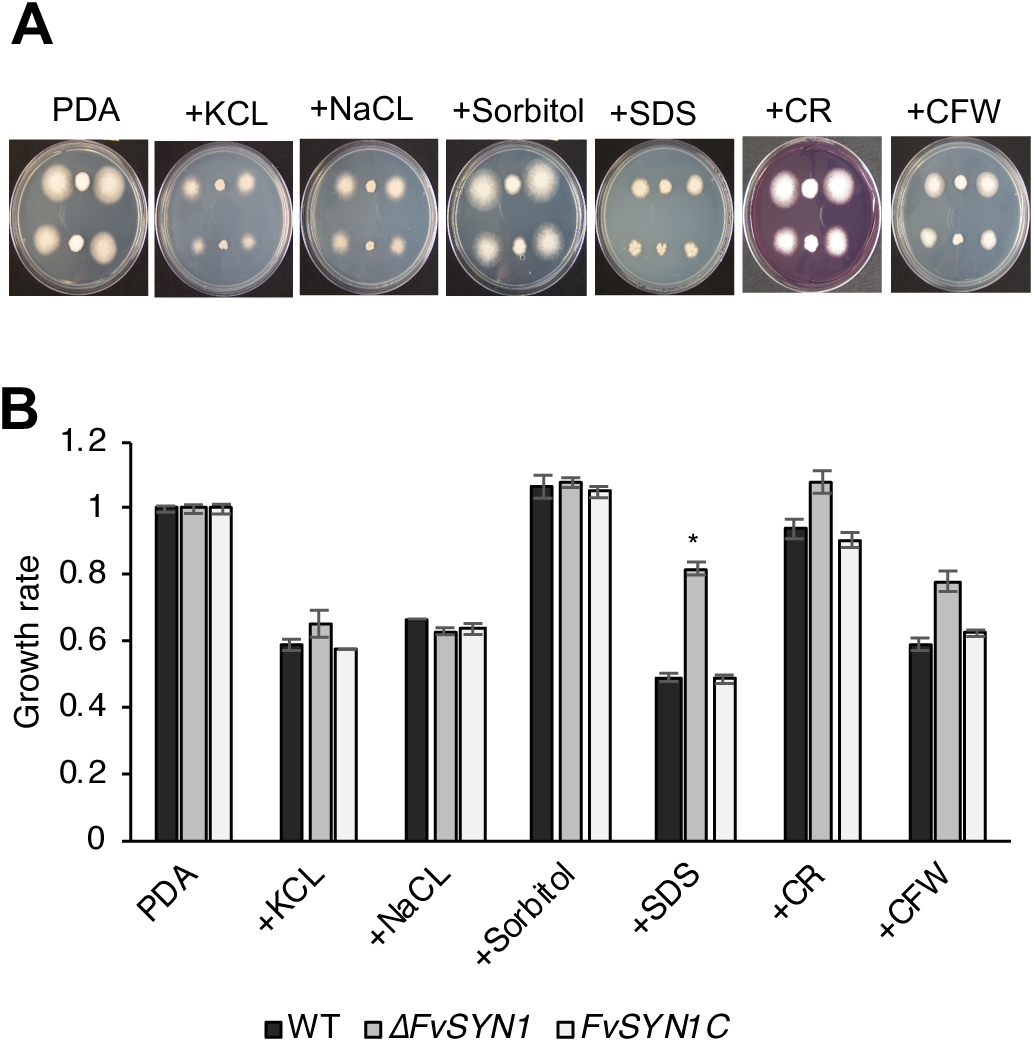
Defects of the Δfvsyn1 mutant in response to various stressors. (A) 4 μl-10^5^ spore suspensions (first row on plates) and 10^4^ spore suspensions (second row on plates) of wild type, mutant and complemented strains were cultured on PDA medium supplemented with hyperosmotic, oxidative stressors and cell wall antagonists for 5 days. SDS, sodium dodecylsulfate; CR, Congo red; CFW, Calcofluor white. (B) The growth rates (artificially set to 1 on PDA plates) of the indicated strains (10^5^ inoculations) under various stress conditions. Error bars represent the standard deviation from three independent experiments and asterisks indicate statistically significant differences (P < 0.05) analyzed by t-Test.

### 3.5 SynN and SNARE domains play different roles in vegetative growth and conidiation

FvSyn1 contains two well-recognized domains, a SynN domain at N-terminus and a SNARE motif at C-terminus. We generated Δ*SynN* (WT gene with a complete deletion of the *FvSYN1* N-terminus with its native promoter and terminator) and Δ*SNARE* (WT gene with a complete deletion of the C-terminus with its native promoter and terminator) mutants to study domain functions (Fig. 5A). Complemented strain FvSYN1C (with wild-type *FvSYN1* gene driven by its native promoter transformed into Δfvsyn1) (Kim *et al*., 2018b) was used as a positive control. When we compared the vegetative growth of these mutants on PDA and V8 plates, Δ*SynN* and Δ*SNARE* mutants exhibited similar growth rate as the Δfvsyn1 mutant, but all three mutants showed significantly slower growth rate than the wild-type progenitor and FvSYN1C (Fig. 5B). Interestingly, the mutants Δ*SynN* and Δfvsyn1 exhibited highly dense and fluffier mycelial growth when compared against all strains. Meanwhile Δ*SNARE* mutant mycelial growth was considered in-between these two mutants and WT when grown on PDA plates (Fig. 5C). When all these strains were inoculated on cracked maize kernels, we did not observe statistical difference in ergosterol production (Fig. 5D). Our earlier study also showed that WT, Δfvsyn1 and FvSYN1C produced similar amount of fungal mass when cultures were harvested from YEPD broth (Kim *et al*., 2018b). When we measured conidia production in these strains, we surprisingly learned that Δ*SynN* mutant produced macrospores while all other strains failed to produce macrospores, which is a deviation from a typical *F. verticillioides* asexual reproduction process under laboratory conditions (Fig. 5E). Our data showed that both domains, SynN and SNARE, are both important for normal growth rate. Meanwhile, SNARE domain is more important for aerial mycelial development while SynN domain is necessary for maintaining normal microconidia production.

**Fig. 5.**
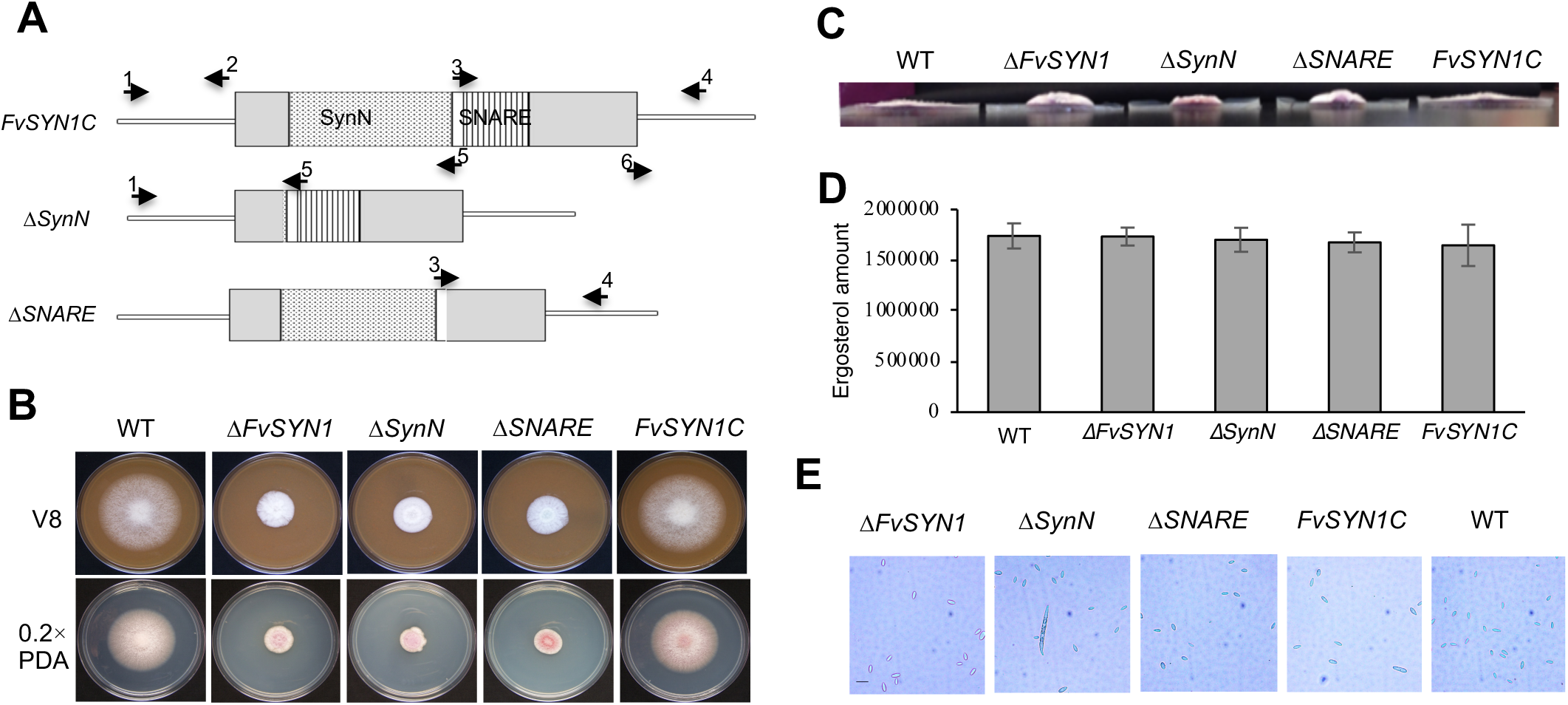
SynN domain and SNARE domain of *FvSYN1* are important for vegetative growth. (A) *FvSYN1* complementation constructs of ΔSynN and ΔSNARE used to test the functional roles of the C-terminus and N-terminus regions. (B)Vegetative growth of WT, *ΔFvSYN1*, *ΔSynN*, *ΔSNARE* and complementation (*FvSYN1C*) strains were examined on V8 and 0.2XPDA agar plates. Strains were point inoculated with a 5μl-10^5^ spore suspensions and incubated for 6 days at 25 °C under 14 h light/10 h dark cycle. (C) A 5μl-10^5^ spore suspensions of WT, *ΔFvSYN1*, *ΔSynN*, *ΔSNARE* and *FvSYN1C* strains were inoculated and incubated on PDA plates for 6 days at 25 °C under 14 h light/10 h dark cycle. Pictures for colony thickness were taken from a side view. (D) Quantification of ergosterol production in *F. verticillioides* strains. 2×10^6^ spores of WT, *ΔFvSYN1*, *ΔSynN*, *ΔSNRE* and *FvSYN1C* strains were inoculated on nonviable autoclaved maize kernels and incubated for 7 days at 25 °C under a 14-h light/10-h dark cycle. Ergosterol production was quantified by high-performance liquid chromatography (HPLC) analysis. All values represent the means of three biological replications with standard errors shown as error bars. (E) WT, *ΔFvSYN1*, *ΔSynN*, *ΔSNARE* and *FvSYN1C* strains were inoculated on V8 plates and spores were collected one week after incubation. Spores were observed under microscope. Scale bar = 20 μm.

### 3.6 SynN domain and SNARE domain in FvSyn1 are important for virulence

To test virulence, spore suspensions of wild-type, Δfvsyn1, Δ*SynN*, Δ*SNARE*, FvSYN1C strains and water (negative control) were inoculated into silver queen maize seedling mesocotyls as described previously (Christensen *et al* 2014; Zhang *et al*., 2018). Seedling rot symptoms were monitored two weeks after incubation. Seedlings inoculated with Δfvsyn1, Δ*SNARE*, and Δ*SynN* mutants showed significantly reduced levels of rot, approximately 60%, 50% and 40% reduction respectively by analyzing average mesocotyl rot symptom area, when compared with the wild-type progenitor. Meanwhile, FvSYN1C complementation strain showed complete restoration of virulence in maize seedlings (Fig. 6A). These results suggested that these two domains in FvSyn1 collectively play a role in *F. verticillioides* virulence.

**Fig. 6.**
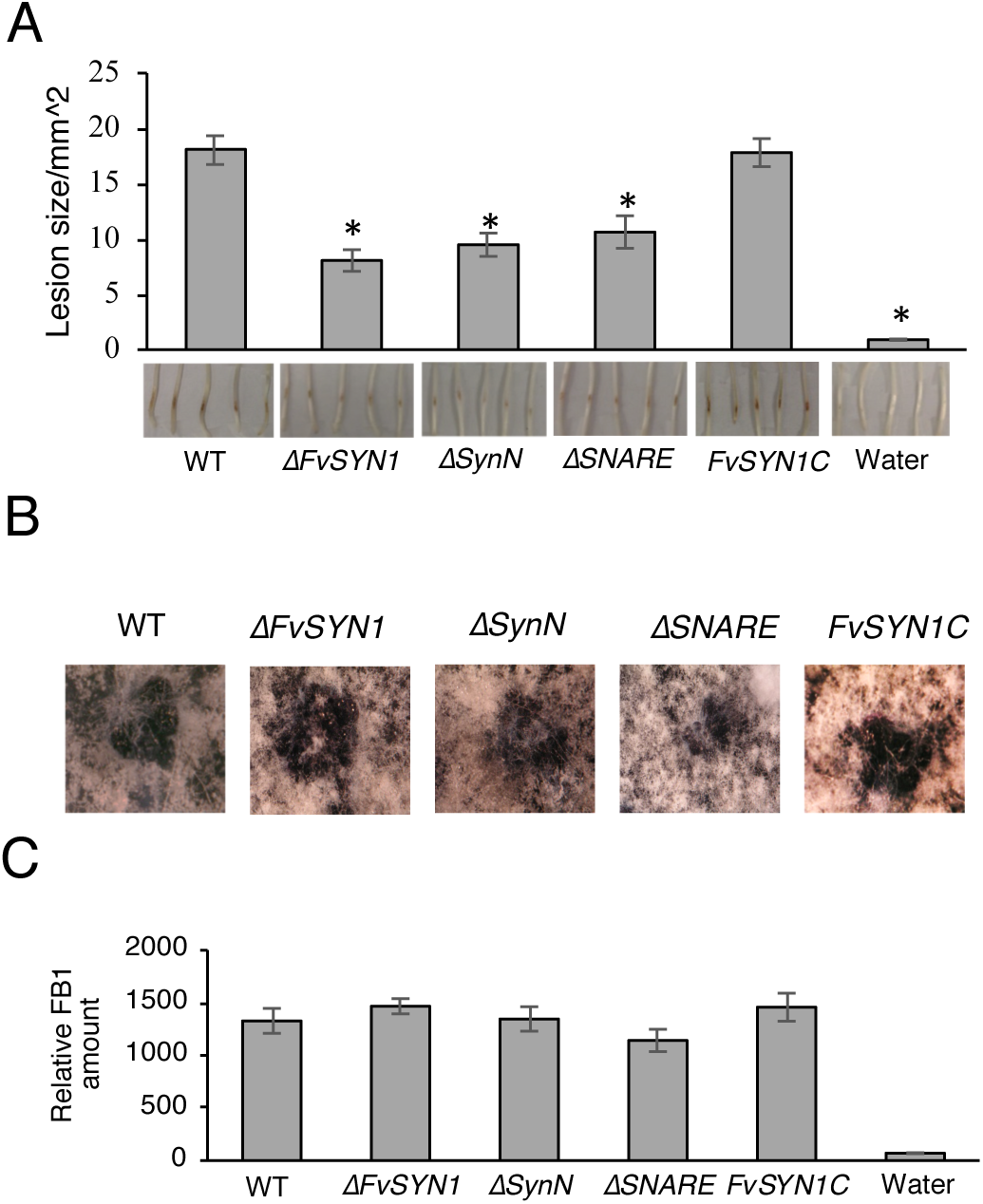
SynN domain and SNARE domain of *FvSYN1* are important for virulence but not dispensable for sexual mating and FB1 production. (A) One week old silver queen maize seedlings were inoculated with 10^7^ /ml spore suspension of fungal strains (WT, *ΔFvSYN1*, *ΔSynN*, *ΔSNARE* and *FvSYN1C* strains) on mesocotyls. Lesion areas were quantified by Image J after 2-week incubation. All values represent the means of three biological replications with standard errors shown as error bars. A sterisk above the column indicates statistically significant difference (P<0.05) analyzed by t-Test. (B) *F. verticillioides* sexual mating assay. Crosses were performed as described previously by Sagaram *et al*. (2007), carrot agar plates were maintained at 25 °C until perithecia and ascospores were observed and characterized. (C) Quantification of FB1production in *F. verticillioides* strains. 2×10^6^ spores of WT, *ΔFvSYN1*, *ΔSynN*, *ΔSNARE* and *FvSYN1C* strains were inoculated on nonviable autoclaved maize kernels and incubated for 7 days at 25 °C under a 14-h light/10-h dark cycle. Ergosterol production was quantified by high-performance liquid chromatography (HPLC) analysis. FB1 biosynthesis was normalized to growth with ergosterol contents. All values represent the means of three biological replications with standard errors shown as error bars.

### 3.7 SynN domain and SNARE domain of FvSYN1 are dispensable for sexual mating and FB1 production

Upon testing *FvSYN1* gene and its key domains for virulence, we questioned how fumonisin production would be impacted by these domains. WT, Δfvsyn1, FvSYN1C strains as well as Δ*SNARE*, Δ*SynN* mutants were inoculated on cracked maize kernels. FB1 levels and fungal biomass were measured after 7 days of incubation, and we normalized FB1 production to fungal growth in all samples tested. *ΔFvSYN1*, *ΔSynN*, *ΔSNARE* and FvSYN1C strains all produced similar level of FB1 to that of the WT strain (Fig. 6B). Therefore, we concluded that *FvSYN1* does not play a role in regulating FB1 biosynthesis. To test whether this gene and its domains are important for sexual mating, Δfvsyn1, Δ*SynN*, Δ*SNARE* mutants as well as WT and FvSYN1C strains were crossed with the opposite mating type *F. verticillioides* strain on carrot agar plates following the standard laboratory method. After 21 days of incubation, all mating pairs resulted in a similar number of perithecia (Fig. 6C) with viable ascospores, suggesting that *FvSYN1* is not critical for sexual reproduction in *F. verticillioides*.

### 3.8 SynN domain and SNARE domain of FvSYN1 are involved in carbon metabolism

To test whether *FvSYN1* gene and its key domains are important for catabolism of carbon sources, WT, Δfvsyn1, Δ*SNARE*, Δ*SynN*, FvSYN1C were inoculated onto media with sucrose, sorbitol, starch or pectin as the carbon source (Fig. 7A). As the carbon source can inhibit or promote the fungal growth, the growth of the strains on the medium with sucrose was set as 1(Fig. 7B and Fig. 7C). The growth of all strains was inhibited on the media supplemented with sorbitol when compared with the growth on the medium with sucrose. However, on the media supplemented with starch and pectin, the growth of WT and complementary strain FvSYN1C were inhibited, while the growth of mutant Δfvsyn1 as well as the domain deletion mutants Δ*SNARE* and Δ*SynN* were enhanced. Taken together, these results suggest that FvSyn1 regulates metabolization of different carbon sources.

**Fig. 7.**
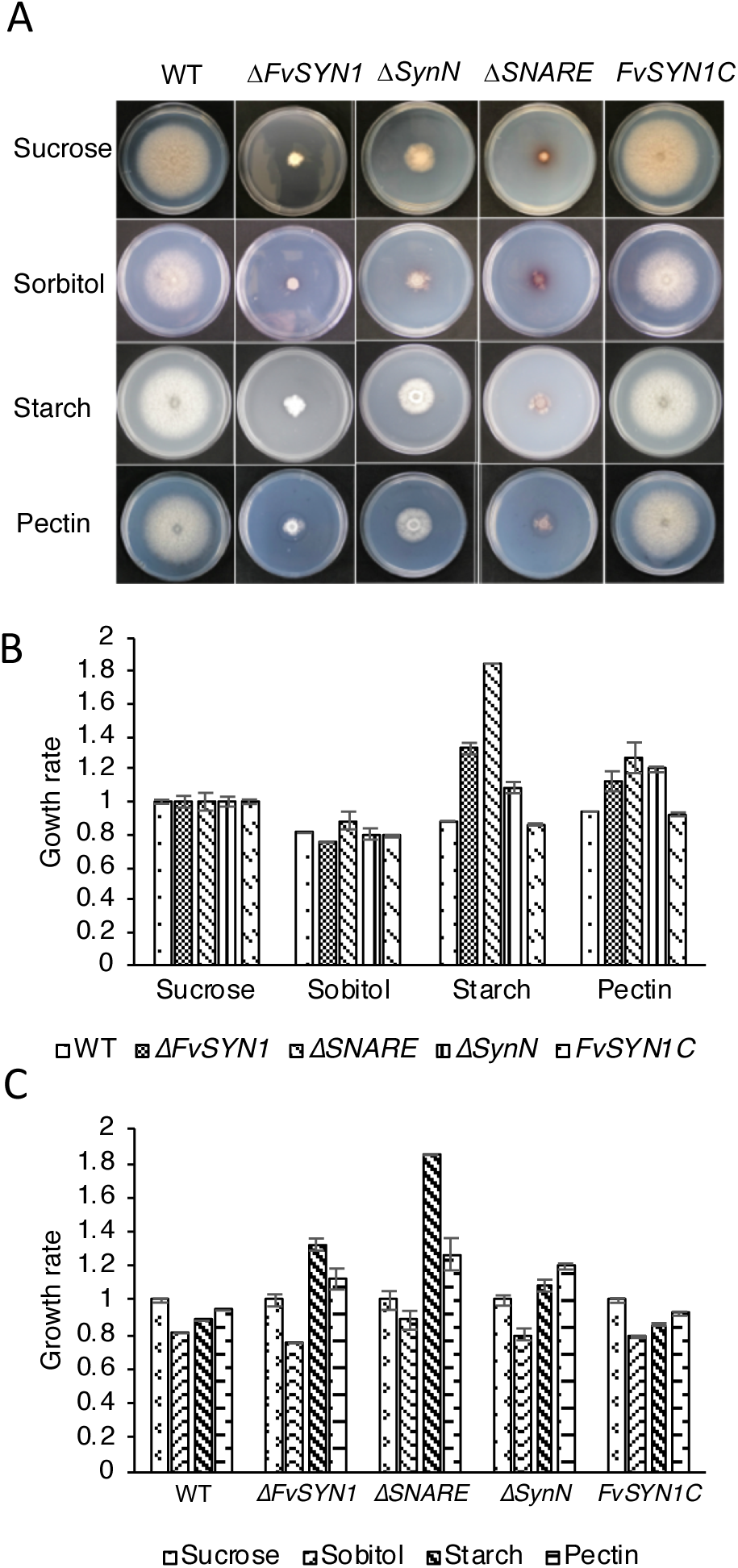
SynN domain and SNARE domain of FvSYN1 are involved in the secretion of catabolism of carbon sources. (A) Spores (4μl of 2 × 10^6^) of all strains were inoculated on media supplemented with different carbon sources, including sucrose, sorbitol, starch, and pectin. (B) Colony diameters at 8 days after inoculation are shown in each panel with standard deviations.

## 4. Discussion

SNARE proteins are well conserved in eukaryotes and play important roles in membrane fusion. In fungi, SNARE proteins in *N. crassa* and *F. graminearum* were shown to be important for hyphal growth, conidiation, fertility and virulence (Gupta *et al*., 2013; Hong *et al*., 2010). In our study, we have shown that the deletion of *FvSYN1* gene resulted in altered hyphal morphogenesis and reduced virulence on maize seedlings. Although there was negative impact on growth when Δfvsyn1 strain was grown on PDA and V8 agar plates, there was no significant difference in fungal mass production when the mutant was grown in liquid broth or in maize cracked kernels. These outcomes suggested that the reduced virulence observed in the mutant may be related to the aberrant hyphal morphology rather than growth deficiency on agar media. Based on literature, we hypothesize that the abnormal hyphal development in FvSyn1 deletion mutant negatively impacts membrane fusion and ultimately delivery of virulence factors important for host infection and colonization.

Studies have shown that SNARE proteins MoVam7 in *M. oryzae* and FgVam7 in *F. graminearum* are important for cell wall integrity, and mutation in these genes resulted in sensitivity against cell wall stressors (Dou *et al*., 2011; Zhang *et al*., 2016). In our study, Δfvsyn1 mutant also exhibited different levels of sensitivity to three cell wall stressors tested. Δfvsyn1 grew better in medium amended with cell wall stressors when compared with wild type and FvSYN1C strains. It has been reported that CFW and CR perturbs cell wall assembly by binding to chitin. SDS is known to act indirectly by perturbation of plasma membrane, resulting in cell wall debilitation and triggering stress responses (Ram and Klis, 2006; Carbó and Pérez-Martín, 2010; Pérez-Nadales and Di, 2016). Based on literature, it is possible that Δ*fvsyn1* mutation alters cell wall remodeling response in *F. verticillioides* by increasing a set of wall maintenance and repair proteins in mutant cell wall, such as increased deposition of chitin in the cell wall. It has been reported that a mutation in *Aspergillus fumigatus fks1* which disturbed the synthesis of β-1,3-glucan resulted in higher chitin content than wild-type cells (Gardiner *et al*., 2005). It could be possible that higher chitin level in Δfvsyn1 mutant causes altered cell wall stress response and less sensitivity to CFW and CR, thus Δfvsyn1 mutant grew better in medium amended with cell wall stressors.

It has been shown that cell wall degrading enzyme pectinase in *F. oxysporum* (García-Maceira *et al*., 2001) and *Botrytis cinerea* (Have *et al*., 1998) play important roles in fungal virulence and host infection. The localization of FvSyn1 on the vacuoles, septa and plasma membrane perhaps can help us explain how abnormal hyphal morphology is influencing the mutant. We hypothesized that defects in protein secretion could lead to reduced virulence in the mutant strain if *F. verticillioides* FvSyn1 plays an important function in the delivery of cell wall degrading enzymes. In the carbon source metabolism assay, we found that FvSyn1 is necessary for proper catabolism of pectin, starch and sorbitol. However, the mutant did not show significant differences in utilization of diverse carbon sources including cellulose, xylan, wheat bran, starch, sucrose and corn meal. Cell wall degrading enzyme activities were tested with 3,5-dinitrosalicylic acid (DNS)-based enzyme assay (Hilton *et al*., 2017), and did not show difference (Fig. S1). While other important enzymes or factors may affect the virulence in the mutants, the mechanism remains unclear and awaits future investigation.

In our study, we observed more fluffier and hyper-branched hyphae phenotype in the Δ*SNARE* mutant, whereas Δ*SynN* mutant exhibited a less fluffy colony growth on agar plates. C-terminal SNARE domain is believed to be directly involved in membrane fusion (Chen & Scheller, 2001; Hanson *et al*., 1997; Lin & Scheller, 1997; Malsam *et al*., 2008). Therefore, our premise was that *F. verticillioides* Δ*SNARE* mutant has a malfunction in membrane fusion, which leads to more fluffier and hyper-branched hyphae phenotype in Δ*SNARE* mutant. Our study also showed that Δ*SynN* mutation resulted in macrospore production, while Δ*FvSYN1* and Δ*SNARE* mutants produced only regular microconidia. It is reported that C-terminal SNARE domain is known to participate in the formation of the core complex (Kee *et al*., 1995). N-terminal domain plays an important regulatory role for SNARE complex assembly, N-terminal domain inhibits complex formation by interacting with the C-terminal SNARE-binding domain (Misura *et al*., 2000; Nicholson *et al*., 1998; Parlati *et al*., 1999, Fernandez *et al*., 1998; Nicholson *et al*., 1998). Based on literature, it is reasonable to predict that Δ*SynN* mutant lost the regulating role on SNARE complex assembly, resulting in macrospore production.

Taken together, our results showed that FvSyn1 performs important roles in hyphal growth, localization, cell wall stress response and pathogenicity in *F. verticillioides*. In order to gain further insight into the virulence mechanism associated with FvSyn1, there is a need to identify and characterize the putative targets of FvSyn1, interacting complex, and involved secretion pathway. One of the key challenges will be whether we can demonstrate how these molecular components associate or coregulate during pathogenesis.

## Acknowledgements

This research was supported in part by the Agriculture and Food Research Initiative Competitive Grants Program Grant (2013-68004-20359) from the USDA National Institute of Food and Agriculture. The authors declare no conflict of interest.

