## Supplementary material for "*Fusarium verticillioides* SNARE protein FvSyn1 harbors two key functional motifs that play selective roles in fungal development and virulence"

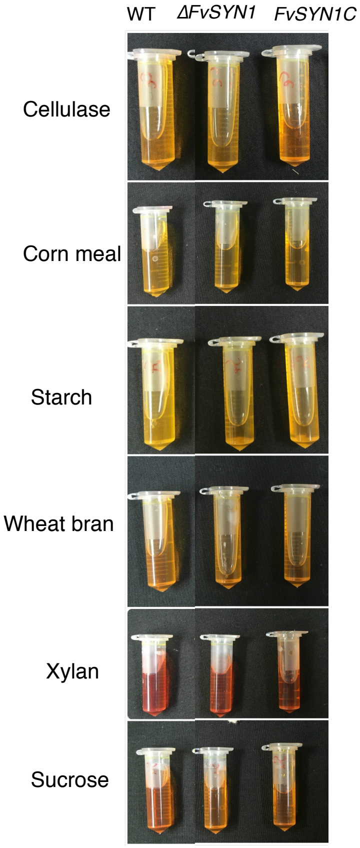


**Fig. S1** Cell wall degrading enzyme activity. Cell wall degrading enzyme activity was tested on diverse carbon sources including cellulose, xylan, wheat byran, starch, sucrose and corn meal as described by Hilton et al (2017).

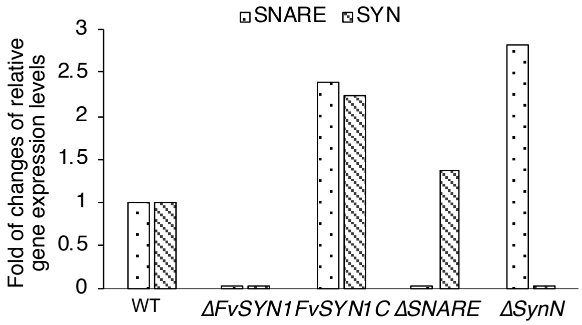


**Fig. S2** qPCR was performed to confirm the motif deletion mutants. SNARE domain primers (Snare-F: GATCTGCTTGGGTATTTGTGTTGC and Snare-R: CCAATGGCTGCCTCGATACTT) and SynN domain primers (SynN-F: AGGGTTATGGAGGTAGCGTTGAG and Synn-R: GATCATGCGGAGCTGTTCCAAG) were used. There is no SNARE domain expression in ΔSNARE mutant when used SNARE domain primers. There is no SynN domain expression in ΔSynN mutant when used SynN domain primers.

**Table S1**. Primers used in this study. The underlined sequences were for fusion purpose.

| **Primer Name** | **5'-3' sequence** |
|---|---|
| 1 | ACC AGG ATT ATT GGA GAG CAG G |
| 2 | TGA GCT GGA GCT CTG CTT TG |
| 3 | A AGC AGA GCT CCA GCT CA CCA GCT CGA CTC GCT TTC C |
| 4 | TTC AAT CCT TGC AGC TGG TG |
| 5 | GGA AAG CGA GTC GAG CTG G |
| 6 | CA GCT CGA CTC GCT TTC C TACCTCAGCTACCGCGGAAC |
| Syn1-GFP-F | GGA GCT GGT GCA GGC GCT GGA GCC GGT GCC ATG AGT GTT AGT TTG CGC CTC C |
| Syn1-GFP-R | TTC AAT CCT TGC AGC TGG TG |
| sGFP/F | AGG AAC CCA ATC TTC AAA ATGGTGAGCAAGGGCGAG |
| 5GAsGFP/Rbs | GGCACCGGCTCCAGCGCCTGCACCAGCTCCCTTGTACAGCTCGTCCATGC |
| RP27-F | ACT ATA GGG CGA ATT GGG TAC TC |
| RP27-R | TTT GAA GAT TGG GTT CCT ACG |
